# Polymorphisms in rs2069845 are associated with IL-6 and soluble IL-6 receptor levels during total joint replacement

**DOI:** 10.1101/2024.10.18.619020

**Authors:** Kyle D. Anderson, Bryan Dulion, John Wong, Niyati Patel, Anne Debenedetti, Craig J. Della Valle, Ryan D. Ross

## Abstract

As the number of patients undergoing total joint replacement (TJR) surgery increases, so does the number of revision surgeries. One driver of implant failure and subsequent revision surgery is peri-implant osteolysis, which is driven by inflammation-mediated bone loss. IL-6 is an inflammatory marker that is elevated during the peri-operative period. Early elevations in IL-6 levels have been linked to osteolysis development. The current study asked whether there is genetic contribution to the IL-6-related peri-operative inflammatory reaction to TJR surgery. Patients undergoing primary TJR (total hip or total knee) provided pre-operative and post-operative blood samples for measurement of the circulating levels of IL-6 and the soluble IL-6 receptor (sIL-6r) and evaluation of allele status of three single nucleotide polymorphisms (SNPs) linked to IL-6 or sIL-6r levels - rs2069845, rs228145, and rs4537545. Circulating sIL-6r levels were associated with allele status in the rs2228145 SNP. More interestingly, allele status in the rs2069845 SNP was associated with the change in circulating IL-6 levels following TJR surgery. Specifically, patients with the A,A allele had increasing levels of IL-6, while those harboring the G,A allele had decreasing levels of IL-6. Although the study was not designed to follow implant survival, due to the critical role of IL-6 in driving peri-implant osteolysis development, the results may suggest that the rs2069845 allele has a role in the success of orthopedic implants. rs2069845 polymorphisms may be a useful patient-specific marker of inflammatory response to TJR surgery.

## INTRODUCTION

The number of revision total hip and knee replacement surgeries performed in the U.S. is projected to grow to between 200,000 and 300,000 by the year 2030 [1]. The most common indications for reversion surgery are implant instability and aseptic loosening [2], which are often attributed to the biological loss of implant fixation caused by particle-induced peri-implant osteolysis [3, 4]. Although advances in materials, particularly the wide-spread use of ultra-high molecular weight polyethylene (UHMWPE) liners have reduced wear rates [5], recent analysis of large-scale joint replacement registries still identify aseptic loosening as the most common cause for total hip revisions and the second most common for total knee revisions [6]. Osteolysis progresses for years prior to diagnosis resulting in substantial bone loss, which makes revision surgeries more challenging [7, 8]. Identifying patients at risk for aseptic loosening may allow for early intervention to halt or reverse bone loss, as has been demonstrated in preclinical model systems and clinical case-reports [9, 10]. However, to date, no validated biomarkers have been established to diagnose osteolysis or subsequent aseptic loosening [11].

The pathophysiology of osteolysis involves inflammation-induced bone loss caused by wear particles generated from prosthetic materials, such as polyethylene [12, 13]. Therefore, it is not surprising that several cytokines have been proposed as late-stage biomarkers of implant failure caused by osteolysis [11]. Recently, our lab has reported that elevated levels of interleukin 6 (IL-6) were among a panel of biomarkers able to prospectively discriminate total joint recipients that would eventually develop radiographically confirmed osteolysis [14]. Perhaps more interestingly, pre-operative IL-6 levels (circulating levels measured prior to implant placement) were equally able to distinguish the patients that would eventually develop radiographic osteolysis, suggesting that patient intrinsic factors exist that predispose patients to osteolysis development and these factors are likely related to elevated IL-6 signaling.

IL-6 is an inflammatory cytokine that induces osteoclastogenesis, leading to bone loss [15, 16]. IL-6 can promote osteoclastogenesis both directly, by stimulating osteoclast progenitor cells, and indirectly via osteoblasts (reviewed here [17]). In osteoblasts, the IL-6 signal is mediated through the interaction of soluble IL-6 with a two-component receptor system. The IL-6 binding partner is the IL-6 receptor (IL-6R) protein, which can exist as membrane-bound protein or as a soluble agonistic molecule (sIL-6R) [18]. The IL-6/sIL-6R complex associates with the membrane-bound gp130 protein to initialize a signaling cascade to increase the production of RANKL by osteoblasts. RANKL will bind to the RANK receptor, expressed by osteoblast progenitor cells, which triggers osteoclastogenesis. Therefore, IL-6 likely serves as an important regulator of osteoclast-mediated bone loss seen with the progression of osteolysis. Indeed, IL-6 is elevated in the peri-implant environment of failed orthopedic implants [19, 20]. Further, single nucleotide polymorphisms (SNPs) in the IL-6 promotor region have been associated with implant lifespan [21, 22] and severity of osteolysis [23], in support of our proposed patient intrinsic factors. The current study aimed to evaluate whether SNP polymorphisms implicated in the regulation of the IL-6 signaling pathway are associated with the early inflammatory response to total hip and knee replacement surgery. We focused on three SNPs previously linked to IL-6/sIL-6r regulation. Specifically, rs2069845 has been implicated in the severity and time to onset of inflammatory reactions in leprosy patients [24], rs2228145 (also referred to as rs8192284) has been linked to sIL-6r levels [25] and coronary heart disease risk [26], and rs4537545 has been implicated in the regulation of circulating IL-6 and sIL-6r levels [27-29]. We hypothesized that allele variance in these SNPs would be associated with changes in circulating IL-6 and sIL-6R in the peri-operative period of patients undergoing primary total joint replacement.

## MATERIALS AND METHODS

The study design was approved by the institutional review board on human research. A signed consent form was obtained from each patient providing permission for storage of biofluids for future research studies.

### A Priori Power Analysis

Prior to initiating patient enrollment, a power analysis was performed to determine the predicted sample size necessary to detect a significant association between the pre- and post-operative levels of IL-6 and sIL-6R and SNP isotypes. The frequency of allele variations in the four proposed SNPs ranged between 32-45% of the population (NCBI 1000 Genomes Browser). Power analysis was performed using the Quanto power calculation tool (http://biostats.usc.edu/software). To power the current study, we used the allele frequencies, the odds ratios we previously determined for pre- and post-operative IL-6 levels to predict osteolysis [14], and an estimated population risk of 1%, which predicts that roughly 23 patients would be necessary. Recruitment was initiated until the target 23 patients were consented and both pre and post-operative blood samples were collected.

### Participant Enrollment

A total of 34 patients receiving either primary total hip or primary total knee replacement surgeries with no prior history of inflammatory disease were consented at a pre-operative consultation to enroll in the present study. Enrollment occurred between March 2019 and August 2022. Participants were provided informed consent forms, which were signed in the presence of the lead investigators – KDA or RDR. A total of 7 participants subsequently canceled their scheduled surgeries and an additional 4 participants were excluded due to missing pre- or post-operative samples. In total, 23 patients receiving primary total hip or knee replacements were included, thereby achieving our sample size.

Blood samples were collected by trained phlebotomists as part of the venipuncture service of the Rush Medical Laboratories. The average time between the pre-operative sample collection date and surgery was 10 (± 10) days and the average time between the post-operative collection data and surgery was 33 (± 16) days. Samples were collected in K2 EDTA blood collection tubes (BD) and processed to obtain plasma and buffy coats by centrifugation at 2,500 RPM for 15 minutes. The resulting phases were aliquoted into fresh tubes and stored at -80ºC.

### Cytokine Quantification

Plasma samples were thawed to room temperature prior to analysis. The circulating levels of IL-6 and sIL-6R were assessed using commercially available ELISAs (Human IL-6 ELISA, BioLegend) (Human IL-6 receptor [soluble] Human ELISA Kit, ThermoFisher). Due to limited plasma volume obtained for some participants, this study could not obtain results for six samples of IL-6 (four pre-operative and two post-operative samples) and one IL-6R (post-operative sample).

### Single Polymorphism (SNP) Isotyping

SNP allele status was assessed using qPCR (Applied Biosciences). Validated Taqman primers for the SNPs of interest – rs2069845, rs2228145, and rs4537545 - were purchased from ThermoFisher. PCR reactions were set up on a 96 well plate and all patient samples were tested in triplicate.

### Statistical Analysis

Prism (Version 8; GraphPad) and SPSS (Version 19.0; SPSS Inc.) software packages were used for plotting and data analysis, respectively. Due to low sample volumes, a total of 6 IL-6 measurements were missing – four in the pre-operative time point and two in the post-operative time point – and one sIL-6r measurement in the post-operative time point.

Prior to testing, data was evaluated for normality using the Shapiro-Wilk test. Roughly half of the datasets were not normally distributed and therefore a more conversative non-parametric approach was used to compare means in all subsequent analyses. In all three SNPs evaluated, the rarest of the allele frequencies was significantly underpowered, therefore we focused on comparing the two more common allele frequencies using a non-parametric Mann-Whitney U test. To evaluate level changes pre- and post-operatively, we used mixed-effects ANOVA models. The effects of allele status, time (pre-vs. post-operative) and the interaction of these two terms are presented.

## RESULTS

### Patient Demographics Based on SNP Allele

The total number of patients receiving TKR was 12, while 11 received THR. There were 7 males and 16 females overall, with an average age of 67.6 (±9.1) years at the time of surgery. Participant demographic data according to participant SNP allele status are presented in Table 1.

**Table 1:** Demographics and SNP alleles for patients undergoing primary total joint replacement surgery.

| SNP Allele Status<br>(sample size, %) | Surgery<br>(# THR vs. TKR) | Sex (n, % female) | Age at Surgery<br>(mean, range) |
| --- | --- | --- | --- |
| rs2069845 |  |  |  |
| A,A<br>(11, 47.8%) | 5 THR<br>6 TKR | 5 (45%) | 67.5 (58-84) |
| G,A<br>(10, 43.4%) | 4 THR<br>6 TKR | 9 (90%) | 67.7 (49-86) |
| G,G<br>(2, 8.7%) | 2 THR<br>0 TKR | 1 (50%) | 67.5 (63-72) |
| rs2228145 |  |  |  |
| C,A<br>(13, 56.5%) | 6 THR<br>7 TKR | 4 (31%) | 67.2 (49-84) |
| A,A<br>(8, 34.7%) | 4 THR<br>4 TKR | 6 (75%) | 66.8 (58-86) |
| C,C<br>(2, 8.7%) | 0 THR<br>2 TKR | 1 (50%) | 73 (49-84) |
| rs2228145 |  |  |  |
| C,T<br>(15, 65.2%) | 9 THR<br>6 TKR | 5 (33%) | 66.0 (49-84) |
| C,C<br>(5, 21.7%) | 2 THR<br>3 TKR | 1 (20%) | 69.0 (58-86) |
| T,T<br>(3, 13.0%) | 0 THR<br>3 TKR | 2 (67%) | 73.0 (70-76) |

### Effects of rs2069845 allele status on peri-operative IL-6 and sIL-6r levels

Neither pre-operative nor post-operative IL-6 levels were affected by rs2069845 allele status (Figure 1). However, there was a significant allele by time interaction when comparing the two more common alleles, A,A and G,A, indicating that patients harboring these two alleles react differently to TJR surgery. Specifically, patients with the A,A allele had increasing IL-6 levels post-operatively, while those with the G,A allele had decreasing IL-6 levels. There were no significant allele differences in the pre-operative sIL-6r levels. Patients with the G,A allele had significantly elevated post-operative sIL-6r levels when compared to those with the A,A allele. There were no significant allele, time, or allele by time interactions in the sIL-6r levels.

**Figure 1.**
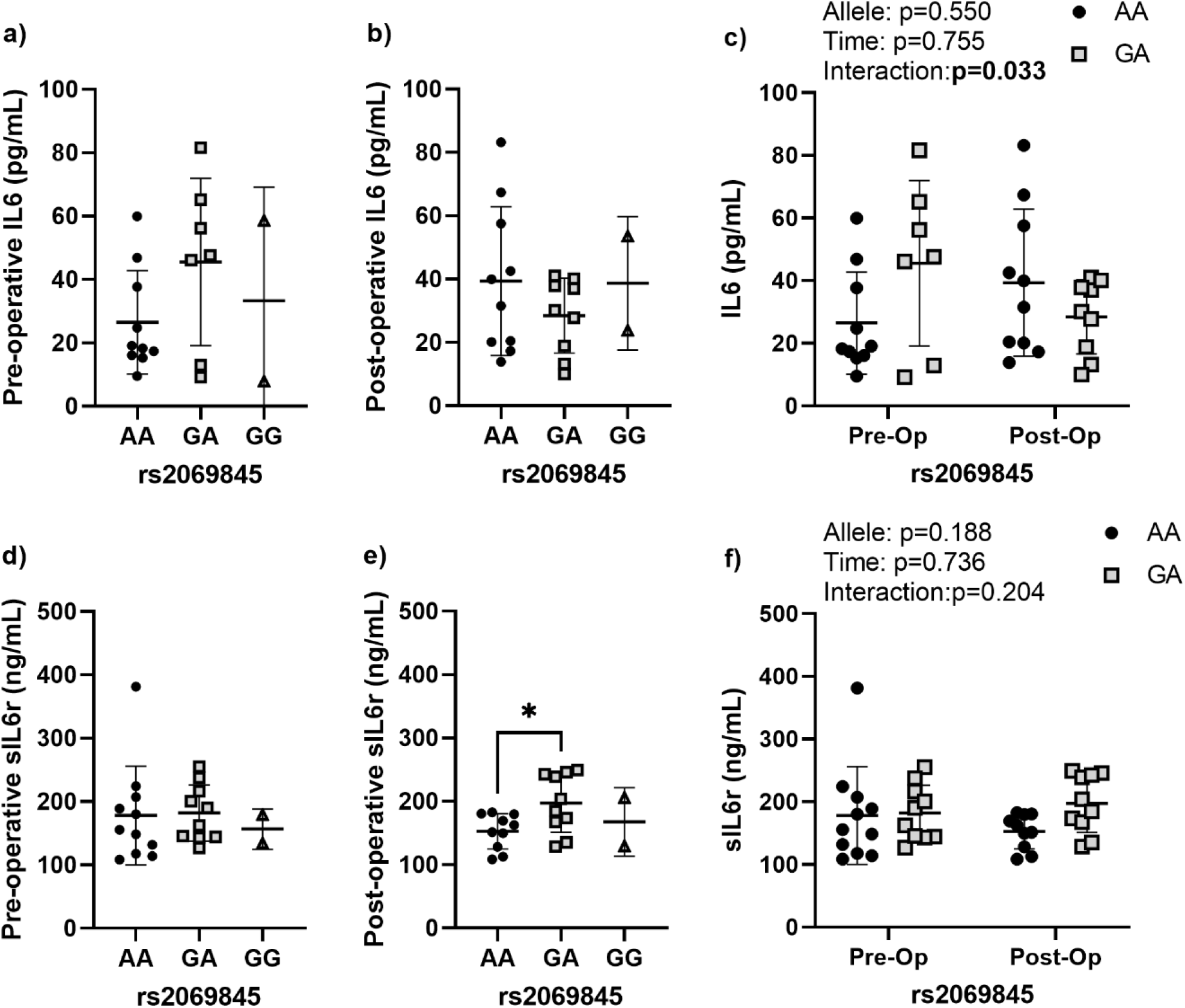
Perioperative IL-6 and sIL-6r dynamics according to rs2069845 allele status. (a) Pre-operative and (b) post-operative circulating IL-6 levels according to allele variant. (c) Comparison of the pre- and post-operative IL-6 levels according to allele variant for the two most prevalent alleles. (d) Pre-operative and (e) post-operative circulating sIL-6r levels according to allele variant. (f) Comparison of the pre- and post-operative sIL-6r levels according to allele variant for the two most prevalent alleles (A,A and G,A). Data are presented as individual measures with the mean and standard deviations. Significant pair-wise comparisons from non-parametric Mann-Whitney U tests are presented, when significant, as bars over the data in panels a, b, d, e. The results from non-parametric mixed effects comparisons of the effects of allele, time, and the allele by time interaction for the two most common alleles are presented as legends in panels c and f.

### Effects of rs2228145 on peri-operative IL-6 and sIL-6r levels

The pre-operative and post-operative IL-6 levels were not affected by rs2228145 allele variant (Figure 2). Nor were there time or interaction effects detected when comparing the pre- and post-operative change. Both pre- and post-operative sIL-6r levels were significantly higher in patients with the C,A allele compared those with the A,A allele, which was confirmed by the significant allele effect noted when comparing both time points together (allele effect: p=0.004).

**Figure 2.**
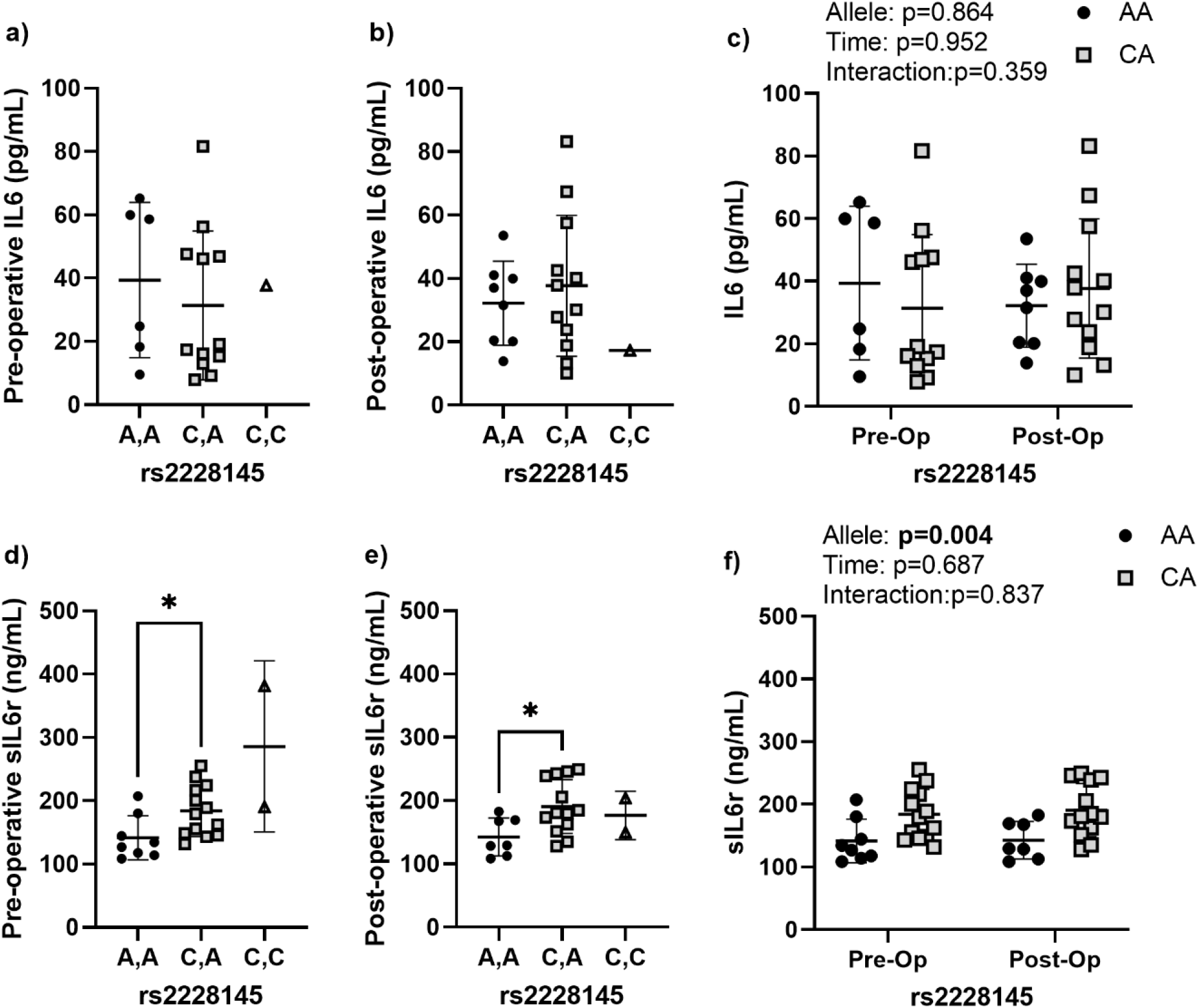
Perioperative IL-6 and sIL-6r dynamics for rs2228145. (a) Pre-operative and (b) post-operative circulating IL-6 levels according to allele variant. (c) Comparison of the pre- and post-operative IL-6 levels according to allele variant for the two most common alleles. (d) Pre-operative and (e) post-operative circulating sIL-6r levels according to allele variant. (f) Comparison of the pre- and post-operative sIL-6r levels according to allele variant for the two most common alleles (A,A and C,A). Data are presented as individual measures with the mean and standard deviations. Significant pair-wise comparisons from non-parametric Mann-Whitney U tests are presented, when significant, as bars over the data in panels a, b, d, e. The results from non-parametric mixed effects comparisons of the effects of allele, time, and the allele by time interaction for the two most common alleles are presented as legends in panels c and f.

### Effects of rs4537545 on peri-operative IL-6 and sIL-6r levels

rs2228145 allele status had near significant effects on both IL-6 and sIL-6r levels (Figure 3). The pre- and post-operative levels of IL-6 were higher in patients with the C,T allele, but the results were not statistically significant, likely due to the relatively few patients with the C,C allele (Figure 3). When both timepoints were compared in the ANOVA, the results showed a trend toward an allele effect (p=0.077). Similar effects were noted in the levels of sIL-6r, which were not significant at either timepoint, but there was a trend for an allele effect (p=0.061).

**Figure 3.**
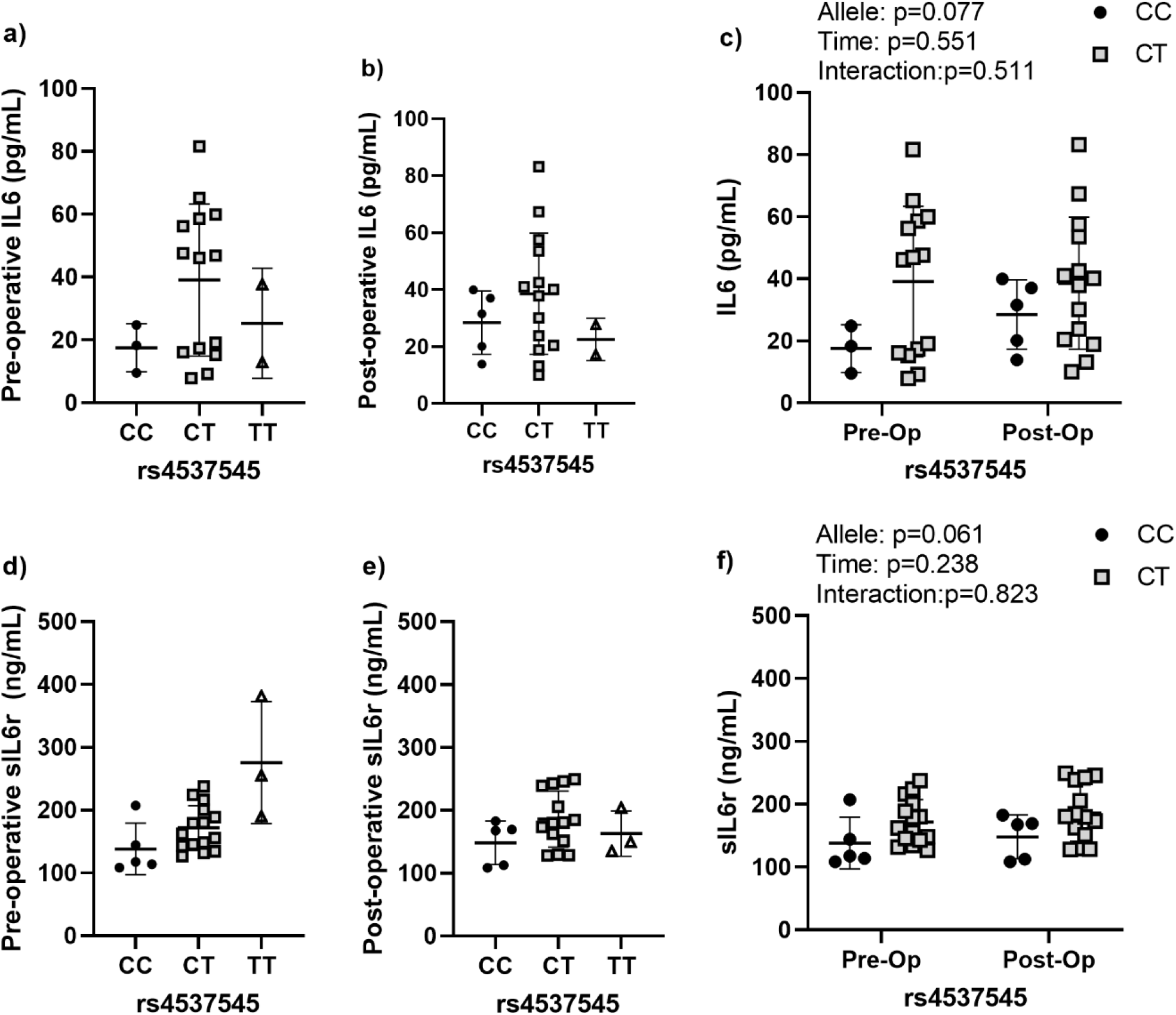
Perioperative IL-6 and sIL-6r dynamics for rs4537545. (a) Pre-operative and (b) post-operative circulating IL-6 levels according to allele variant. (c) Comparison of the pre- and post-operative IL-6 levels according to allele variant for the two most common alleles. (d) Pre-operative and (e) post-operative circulating sIL-6r levels according to allele variant. (f) Comparison of the pre- and post-operative sIL-6r levels according to allele variant for the two most common alleles (C,C and C,T). Data are presented as individual measures with the mean and standard deviations. Significant pair-wise comparisons from non-parametric Mann-Whitney U tests are presented, when significant, as bars over the data in panels a, b, d, e. The results from non-parametric mixed effects comparisons of the effects of allele, time, and the allele by time interaction for the two most common allele are presented as legends in panels c and f.

## DISCUSSION

Peri-implant osteolysis and the subsequent loss of implant stability is one of the primary causes of total joint replacement failure [2-4, 6]. There are currently no treatment options to halt or reverse peri-implant osteolysis progression. Osteolysis is generally only identified when a patient presents with pain or instability, at which point the only corrective action available is revision surgery. Revision surgeries for hip and knee replacements are more complex, costly, and have worse outcomes when compared to primary surgeries [30-32], which are largely driven by the substantial bone loss associated with peri-implant osteolysis. Identifying risk factors for the development of osteolysis may allow for more routine clinical monitoring of at-risk patients, potentially allowing for early intervention prior to substantial bone loss and are critical for the future design of clinical trials for pharmaceutical treatments for osteolysis.

We have previously identified IL-6 levels as an early biomarker of subsequent osteolysis in one of the few longitudinal biomarker studies in total joint replacement patients [14]. Importantly the patients that ultimately developed osteolysis had lower pre-operative IL-6 levels but higher post-operative levels than patients that did not develop osteolysis over the same period, suggesting a possible patient-intrinsic sensitivity to TJR surgery and implant materials. The potential genetic contribution to osteolysis has been summarized by Jagga et al [33], which highlighted a variety of SNPs that have been linked to the biological response to implant wear particles or risk of osteolysis. Many of the SNP alleles that have been linked to TJR outcomes are involved in the regulation of pro-inflammatory cytokines, such as IL-6.

IL-6, a major cytokine that controls acquired immunity and chronic inflammation [34], is elevated in the peri-implant tissues of implants that failed due to osteolysis [19, 35-39]. IL-6 has also been studied as a circulating biomarker of osteolysis with mixed success [40], likely due to the fact that the majority of studies have investigated IL-6 levels at the end stage of osteolysis [41-44], after osteolysis lesions were detected. Longitudinal assessments of IL-6 levels following total knee arthroplasty are reported to increase immediately post-operatively, reaching a peak as early as 48 hours after surgery [45]. These early changes in IL-6 are also reported in a variety of elective surgeries (summarized herein [46]), but prolonged elevation of IL-6 has been associated with adverse post-surgical complications, including mortality [47-49]. In the bone microenvironment, IL-6 induces the osteoblast production of RANKL, which subsequently activates osteoclast-mediated bone resorption [15, 16]. Therefore, early and prolonged elevations in IL-6 levels likely contribute to sustained osteoclast activity, osteolysis development, and the loss of implant fixation.

Several single nucleotide polymorphisms (SNPs) have been reported to control the expression and circulating levels of IL-6. While we did not find any relationships between the evaluated SNPs and the pre- or post-operative levels of circulating IL-6, we note an interesting SNP by time interaction related to rs2069845 allele status. Specifically, patients harboring the A,A allele had the expected increase in IL-6 following TJR surgery, while those with the G,A, allele saw decreasing IL-6 levels post-operatively. Although preliminary, these results tend to suggest that the G,A may protect from inflammation-induced peri-implant bone loss. It is also possible that these findings represent a faster resolution of post-surgical inflammation, as leprosy patients harboring the G,A allele status in rs2069845 had significantly shorter times to reactional episodes [24]. This same study also noted no associations between rs2069845 allele status and the circulating levels of IL-6, which tends to suggest that this SNP is primarily involved in induced inflammatory reactions. Whether rs2069845 is associated with the risk of orthopedic implant failure is currently unknown, but efforts to reduce IL-6 in preclinical models of particle-induced osteolysis have noted significantly reduced osteoclastogenesis following IL-6 antibody treatment [50], suggesting that lower levels of IL-6 post-operatively may be protective.

sIL-6r is a marker of inflammation that potentiates the effects of IL-6 and polymorphisms in sIL-6r have likewise been associated with levels of both sIL-6r and IL-6 itself. Similar to previous publications, we found that rs2228145 was associated with the circulating levels sIL-6r [27, 51, 52]. Specifically, we noted that patients with the C,A allele in the rs2228145 had higher levels of sIL-6r both pre- and post-operatively. Patients with the C,T allele in rs4537545 also had elevated sIL-6r levels pre- and post-operatively but the results were not significant (p=0.077). The C,T allele was also associated with elevated IL-6 levels both pre- and post-operatively, but again the elevation fell short of the significance threshold (p=0.061). Together, these results suggest that variance in rs4537545 and rs2228145 alleles are likely to have higher baseline inflammation, but that their inflammatory environment is largely unaffected by TJR surgery.

Our study is not without limitations. The sample size is relatively limited. While we performed an a-priori power analysis to identify the number of patients necessary to detect a response, we were underpowered to assess the contribution of rarer allele statuses. The data collected within the current study can be used to better power larger studies focused. Further, we do not know whether any of the patients recruited developed peri-implant osteolysis. Patients were recruited between 2018 and 2020 and because few patients ultimately develop osteolysis and those that do only present with osteolysis years after primary surgery, follow-up studies would take years to complete and require much larger sample sizes. However, as we have previously reported that early IL-6 levels are predictive of subsequent osteolysis [14], these data provide potential mechanistic insight linking a patient’s genetics to the early IL-6 response to TJR surgery. Finally, the timing of post-operative sample collection ranged from 15 to 52 days after surgery. The post-operative sample collection occurred during the regularly scheduled post-operative in-person visit and was subject to patient availability. At least two studies have evaluated circulating IL-6 levels longitudinally following TJR surgery and reported that IL-6 levels are back to pre-operative levels two weeks after uncomplicated surgery [45, 53]. Therefore, it is likely that despite the sample collection variability, all patients had recovered to a new baseline following the initial surgery-induced IL-6 spike, which is reported to occur within the first 48 hours.

## Conclusions

Understanding any genetic factors that could predict the inflammatory response to TJR could prove useful in preventing peri-implant osteolysis. In the present study, we determined that a SNP allele status related to the regulation of IL-6 signaling, particularly in the rs2069845 SNP, was related to the inflammatory response to surgery. Due to the critical role of IL-6 in mediated inflammatory-induced bone loss, these results further our understanding of the potential genetic contribution to the development of peri-implant osteolysis and subsequent implant failure.

## ACKNOWLEDGEMENTS

The study was approved by the Rush University Institutional Review Board number protocol number 17061902-IRB01. The research was supported by the Rush Scientific Leadership Council through the Searle Innovators Award to RDR. The authors thank the patients at Midwest Orthopaedics at Rush that volunteered to take part in this study. We also thank Dr. Vasili Karas and his research staff for their assistance in the recruitment of joint replacement patients.

